# The herbicide acetochlor causes lipid peroxidation by inhibition of glutathione peroxidase 4

**DOI:** 10.1101/2023.04.14.536563

**Authors:** Fahmi Mesmar, Maram Muhsen, Jason P. Tourigny, Jason M. Tennessen, Maria Bondesson

## Abstract

Obesity is increasing worldwide, particularly in rural communities, where people are likely exposed to high levels of pesticides. We here investigated whether six commonly used agricultural pesticides on corn and soy fields have adipogenic activity and act as obesogens. Exposure to two pesticides, the herbicides acetochlor and metolachlor, induced adipogenesis *in vitro* in mouse 3T3-L1 preadipocytes. The most potent compound, acetochlor, was selected for further studies in zebrafish. Acetochlor exposure caused morphological malformations and lethality in zebrafish larvae with an EC_50_ of 7.8 µM and an LC_50_ of 12 µM. Acetochlor exposure also resulted in lipid accumulation is zebrafish larvae when simultaneously fed a high cholesterol diet. To decipher the molecular mechanisms behind acetochlor action, we preformed transcriptomic and targeted lipidomic analysis of exposed animals. The combined omics results suggested that acetochlor exposure increased Nrf2 activity in response to reactive oxygen species, as well as induced lipid peroxidation and ferroptosis. We further discovered that acetochlor structurally shares a chloroacetamide group with known inhibitors of glutathione peroxidase 4 (GPX4). Computational docking analysis suggested that acetochlor covalently binds to the active site of GPX4. Consequently, Gpx4 activity was efficiently repressed by acetochlor, and lipid peroxidation was increased in zebrafish. We propose that acetochlor disrupts lipid homeostasis by inhibiting Gpx4, resulting in accumulation of lipid peroxidation, 4-hydroxynonenal, and reactive oxygen species in the cells, which in turn activate Nrf2. Because metolachlor, among other acetanilide herbicides, also contain the chloroacetamide group, inhibition of Gpx4 activity may represent a novel, common molecular initiating event of obesogens.

**Synopsis:** Rural populations have a high prevalence of metabolic disease and are highly exposed to pesticides. This study reports that the herbicide acetochlor, heavily used on soy and corn fields, inhibits an enzyme that protects from oxidation of lipids in the cell membrane, oxidative stress and a type of cell death called ferroptosis, features that are linked to metabolic disruption and obesity.

## Introduction

Despite the extensive public attention paid to obesity and related metabolic disorders, obesity rates have continued to increase worldwide [1]. In the U.S., the prevalence of obesity has reached epidemic levels among both children and adults [2]. However, the incidence of obesity is not uniform across the country, but rather is significantly higher in rural communities when compared to metropolitan areas [3-7]. Because obesity is associated with an increased risk of chronic diseases including hypertension, diabetes, and cardiovascular disease, such high obesity rates impose a disproportionate burden on rural communities [5].

Obesity is a multifactorial disease that stems from a complex interplay between genetics, caloric intake, and physical inactivity. However, mounting evidence suggests that environmental factors, such as exposure to chemical pollutants, can also contribute to obesity and obesity-related diseases (reviewed in [8]). In this regard, the elevated rates of obesity in rural populations raises the question as to whether agricultural pesticides contribute to the prevalence of metabolic disease.

Since 1970, the U.S. has approved more than 500 pesticides for use in agriculture, and billions of pounds of these chemicals are applied to farm fields each year [9]. Yet, despite widespread use, many pesticides have not been adequately assessed for potential adverse effects on human health. We here examined six heavily used pesticides in the State of Indiana. We first investigated the ability of the pesticides to enhance adipocyte differentiation and lipid droplet formation in mouse 3T3-L1 preadipocytes - a well-established *in vitro* system for identifying and studying obesogenic compounds [10].

The most potent compound causing adipogenesis *in vitro*, the herbicide acetochlor, belongs to a group of acetanilide herbicides, which also include metolachlor and alachlor. These compounds are widely used on soy and corn fields and around 60 million pounds/year were applied in the U.S. between 1980 to 2008 [11]. Because of their abundant use, acetochlor, metolachlor and alachlor leak to waterways and have been detected in lake water at concentrations around 0.05, 0.5 and 0.01 µg/L, respectively (0.03 – 1.8 nM, median values 1992-2006) [12]. However, occasionally the concentrations of acetanilide herbicides spike and up to 300 µM metolachlor was detected in the year 2000 [12]. Farmers applying these herbicides are at risk of being exposed to high concentrations; the geometric mean urinary acetochlor and metolachlor metabolite concentrations were 8.0 μg/L (∼30 nM) and 4.7 μg/L (∼11 nM), respectively, in farmers after they had sprayed their fields [13]. After our initial screen *in vitro*, acetochlor was selected for further analysis *in vivo* using zebrafish larvae.

Larval zebrafish is increasingly used for toxicity testing of pesticides, environmental pollutants, and various industrial chemicals [14-17]. As zebrafish larvae provide a whole-animal model with key tissues, organs, and genes that are involved in the complex processes of metabolic control [18, 19], studies of lipid dysfunction and obesity can informatively be performed in zebrafish [20, 21]. Thus, the zebrafish model has been used for screening for and investigation of metabolic disruptors and obesogens [20].

Through multi-omics using a combination of mRNA sequencing and targeted lipidomics, we investigated the mode of action of acetochlor, and subsequent mechanistic studies and computational tools were used to confirm the predictions. Overall, our studies highlight the power of combining *in vitro* cell culture approaches with studies in zebrafish and indicate that a commonly used herbicide functions as an obesogen in mammalian cells and vertebrate animals. We further propose a molecular model for how acetochlor disrupts metabolic homeostasis.

## Materials and methods

### Chemicals

All chemicals were of PESTANAL® analytical or HPLC grade and were purchased as follows: acetochlor, atrazine, dicamba, flumetsulam, glyphosate, metolachlor, predominantly S-metolachlor, rosiglitazone and ML162, all from Millipore Sigma (St. Louis, MO). The stock solutions were prepared in dimethylsulfoxide (DMSO) (Millipore Sigma) except glyphosate, which was prepared in phosphate buffered saline; PBS (pH 7.4). To avoid freeze-thaw cycles, stocks were stored as aliquots at -20 °C until use. The final dilutions of these stock solutions to be used in the cell cultures or zebrafish treatments always resulted in a DMSO concentration below 0.1%. Full chemical names are provided in the Supporting Information, Materials and Methods and Fig. S1.

### 3T3-L1 cell culture, differentiation, and Nile Red/Nuclear Blue staining

Mouse 3T3-L1 cell pre-adipocytes (ATCC, Manassas, VA) were maintained in pre-adipocyte expansion media consisting of Dulbecco’s Modified Eagle Medium – High Glucose; DMEM-HG (Thermo Fisher, Waltham, MA) supplemented with 10% Donor Bovine Serum with Iron (Thermo Fisher) and 1% Antibiotic-Antimycotic; anti-anti (Thermo Fisher) in 5% CO_2_ at 37 °C until 80% confluency. The differentiation of 3T3-L1 cells in the presence and absence of pesticides and Nile Red/Nuclear Blue staining was performed according to our published protocol [22]. Details of the protocol is described in the Supporting Information, Materials and Methods, and schematically depicted in Figure S2A.

### 3T3-L1 ORO staining, imaging, and quantification

The 3T3-L1 cells were seeded into 6-well plates and differentiated for 8 days with and without the compounds (see Supporting Information, Materials and Methods). The media was gently removed, and the cells were rinsed 3-4 times in PBS and fixed overnight in 4% paraformaldehyde (PFA) at 4°C. The cells were washed in 60% isopropanol. Oil Red O (ORO) stain (Millipore Sigma) was used to quantify intracellular lipid droplets. The ready to use ORO stock in 0.5% isopropanol was diluted (3 parts of stock mixed with 2 parts of miliQH_2_O), filtered with 0.45 µm syringe filter, and incubated on the cells for 20 minutes at room temperature. The cells were rinsed with tap water until the plate became clear. 100 tiles were imaged at 10X magnification with a color camera (DMC-4000) using a DMi8 automated microscope (Leica Microsystems Inc., Buffalo Grove, IL). Pixels were quantified using ImageJ. Briefly, images were converted to 8-bit, the thresholds were carefully adjusted, and the particles were counted.

### Zebrafish husbandry

All animal work was approved by the Indiana University, Bloomington Institutional Animal Care and Use Committee. Maintenance of the adult fish in our system is described in [23] and detailed in the Supporting Information, Methods and Materials.

### *In vivo* exposure, lipid accumulation assay, and quantification

Wild type NHGRI-1 fish (Zebrafish International Resource Center (ZIRC) at the University of Oregon, OR) were placed in breeding tanks overnight. The lipid accumulation assay was performed according to our protocol in [24]. Briefly, the fish were exposed to acetochlor from 3 dpf and fed a 0.005% (w/v) high fat diet (HFD; chicken egg yolk powder, Magic Flavors (Seattle, WA) from 6 dpf. At 11 dpf, the larvae were euthanized, and stained with ORO. The ORO staining was performed three times with at least 20 larvae. ORO-stained larvae were imaged at 5X magnification with a color camera using DMi8 automated microscope (Leica Microsystems Inc.). For quantification of lipid accumulation, at least 12 larvae were pooled together per sample in 1.5 ml tube and incubated for 1 h at room temperature in 200 µl of the extraction buffer: 4% IGEPAL (Millipore, Sigma) in 100% isopropanol. The absorbance of extracted ORO stain was measured on a microplate reader (BioTek Synergy H1; Agilent Technologies, Santa Clara, CA) at 495nm [25]. More details on the zebrafish lipid accumulation assay are available in the Supporting Information, Materials and Methods and in Figure S2B.

### RNA extraction, cDNA synthesis, and quantitative real-time qualitative PCR (qPCR)

The 3T3-L1 cells were seeded in 6-well plate and differentiated as described above with the chemical treatments for 8 days. Total RNA was extracted from the cells using TRIzol (Thermo Fisher) and purified using the RNeasy mini kit (Qiagen, Germantown, MD) according to manufacturer’s protocol. 1 µg of RNA was used for cDNA synthesis using High-Capacity RNA-to-cDNA™ Kit (Thermo Fisher) that utilizes both random octamers, and oligo dT-16 for the synthesis reaction. qPCR experiments were run in technical and biological triplicates using PowerUp™ SYBR™ Green Master Mix (Thermo Fisher) in Applied Biosystems QuantStudio™ 5 Real-Time PCR system (Thermo Fisher). Delta-delta threshold cycle ΔΔC_T_ method was applied to calculate relative gene expression. Primer sequences are listed in Table S1, Supporting information. The data was normalized to the rosiglitazone treatment for statistical calculations.

### Zebrafish embryo toxicity test

Wildtype zebrafish embryos were tested according to a protocol adapted from Acute Fish Toxicity (AFT) test (OECD Test Guideline 236) [26]. Wild type NHGRI-1 adult fish were placed in breeding tanks overnight. In the following morning, the embryos were collected in E3 in 10 cm plate and placed in the incubator until they reached the shield stage. At 6 hpf, 12-15 embryos were placed into wells of a 6-well plate containing 5 ml of the exposure media. 12 embryos for each concentration were transferred individually in 200 µl exposure media to each well of a 96-well plate. The plates were then covered with breathable adhesive seals (Thermo Fisher) to prevent evaporation then placed again in the incubator. The exposure media was not renewed during the experiment. The lethality and morphological characteristics of the embryos were assessed at 72 hpf with a Leica DMi8 microscope equipped with a DMC-4000 color camera. We scored the mortality endpoint based on coagulated yolk and lack of heartbeat. The morphological endpoints scored for were lack of hatching, yolk malformation, heart edema, skeletal deformities such as spinal curvature, and body position, which were assessed and scored according to criteria adapted from previously published studies [27-29]. Each experimental group was treated in triplicates and data were pooled to generate dose-response curves.

### RNA extraction, library construction, and sequencing of zebrafish samples

Hatched zebrafish larvae were treated with two different concentrations of acetochlor (0.78 µM or 7.8 µM) from 48 hpf until 72 hpf. The exposed larvae were assessed for abnormalities using stereomicroscope (MZ6; Leica Microsystems Inc.). Only larvae without gross abnormalities were selected for further analysis. Twenty – 30 larvae were pooled for RNA extraction. Samples were extracted using TRIzol and purified using the RNeasy mini kit according to manufacturer’s protocol. The integrity of the RNA was checked with Agilent TapeStation, samples with RIN >7 were processed using the following steps. From each sample 300 ng of total RNA was used for generating polyA selected libraries following the standard protocol of Illumina strand-specific mRNA library preparation kit. The cleaned adapter-ligated libraries were pooled and loaded on the NextSeq 500 (Illumina Inc. San Diego, CA) with a high output 75 cycle flow cell to generate paired-end reads. Sequencing data has been submitted to GEO: Accession number GSE225543.

### Transcriptome assembly, identification of differentially expressed genes, and pathway analysis

RNA-Seq read quality was assessed with FastQC [30] and MultiQC [31]. Reads were pseudo-aligned and quantified using Salmon v1.6.0 [32] and the D. rerio GRCz11 reference assembly, coding and noncoding transcriptomes retrieved through Ensembl (https://www.ensembl.org), using the chromosomal assembly as the index decoy and the default quantification options. Salmon quantifications were imported into DESeq2 [33] for differential gene expression. Cut-off values were selected using log fold change ≥ |0.4| and FDR < 0.05. Networks, functional analyses, and biological pathways were generated using QIAGEN’s Ingenuity Pathway Analysis (IPA) (https://digitalinsights.qiagen.com/IPA).

### Targeted lipidomics

For targeted lipidomics, approximately 25 wildtype zebrafish larvae were exposed to 7.8 µM acetochlor from 48 to 72 hpf. After 24 h exposure, larvae were collected on ice in 1.7 mL Eppendorf tubes, media removed, and then snap frozen in liquid nitrogen. The targeted lipidomics were performed at the Metabolomics Core at the University of Utah School of Medicine Center (Salt Lake City, UT), and the procedures are detailed in the Supporting Information, Materials and Methods. Six replicates each of control and acetochlor exposure was analyzed.

### Measurement of glutathione peroxidase 4 (Gpx4) activity in zebrafish larvae

Approximately 15 hatched zebrafish larvae were treated with 7.8 µM acetochlor or with 2 µM ML162, a GPX4 inhibitor (Millipore Sigma), for 24 h starting from 48 hpf. The pooled larvae were homogenized in 200 µl PBS with 0.05% Tween 20 with a pestle. The suspension was centrifuged at 15,000 x g at 4 °C for 15 minutes. Zebrafish Gpx4 activity was measured of the supernatant according to manufacturer’s protocol (Millipore Sigma). Total protein was measured using DC protein assay (Bio-Rad Laboratories, Hercules, CA). The Gpx4 activity was calculated and is expressed as U/L per mg of total protein.

### Measurement of 4-hydroxynonenal (4-HNE) in zebrafish larvae

Approximately 50 zebrafish larvae were treated with 7.8 µM acetochlor for 4 days starting from 6 hpf. The media and chemical treatments were renewed at 48 hpf. At 96 hpf zebrafish larvae were collected and snap frozen. The pooled larvae were homogenized in 200 µl PBS with 0.05% Tween 20 with a pestle. The suspension was centrifuged at 12,000 x g at 4 °C for 10 minutes. 4-HNE level was determined by OxiSelect HNE adduct competitive ELISA kit according to manufacturer’s protocol (Cell BioLabs Inc. San Diego, CA). 4-HNE levels were normalized to total protein and data are expressed as µg/ml per mg of total protein.

### Covalent docking

The covalent docking was performed according to Molecular Forecaster’s workflow [34-36]. PREPARE and PROCESS programs were used to prepare protein pdb file for docking. CONVERT and SMART programs were used to prepare the ligand 3D-structure and prepare it for docking. FITTED docking program that was used to predict the acetochlor - GPX4 covalent interaction. BIOVIA Discovery Studio Visualizer was used to visualize the interaction (BIOVIA, D.S. (2015) Discovery Studio Modeling Environment; Dassault Syst. Release, San Diego, CA) [37].

### Statistical analysis

GraphPad Prism software (version 9, GraphPad Software Inc.) and R statistical software (RStudio) were used to generate the graphs and statistical analysis. One-Way ANOVA and student’s t-test were used as parametric statistical procedures. Two-Way ANOVA was used to test data with two categorical variables. Nonparametric Kruskal–Wallis test was used to evaluate data that did not fit normality assumption. False discovery rate (FDR) calculations were included in the identification of differentially expressed genes in RNA-seq experiment.

## Results and Discussion

### Pesticide exposure induces adipogenesis in 3T3-L1 cells

We set out to investigate the adipogenic potential of pesticides widely used in the US state of Indiana, including atrazine, metolachlor, acetochlor, glyphosate, dicamba, and flumetsulam (**Fig. S1**, Supporting Information). As a first step towards this goal, we used murine 3T3-L1 preadipocytes in an *in vitro* assay that examines adipogenesis and lipid droplet formation [22] (see **Fig. S2A**, Supporting Information, for experimental design) [38]. Exposures to acetochlor and metolachlor resulted in a significant lipid droplet accumulation in 3T3L1 cells at 1 and 10 µM when assayed by AdipoRed staining **(Fig. 1A and Fig. S3A**, Supporting Information) and at 10 µM when stained by Oil-Red-O (ORO) and quantified by ImageJ (**Fig. 1B and Fig. S3B**, Supporting Information). Acetochlor was the most potent agent inducing adipogenesis followed by metolachlor.

**Figure 1.**
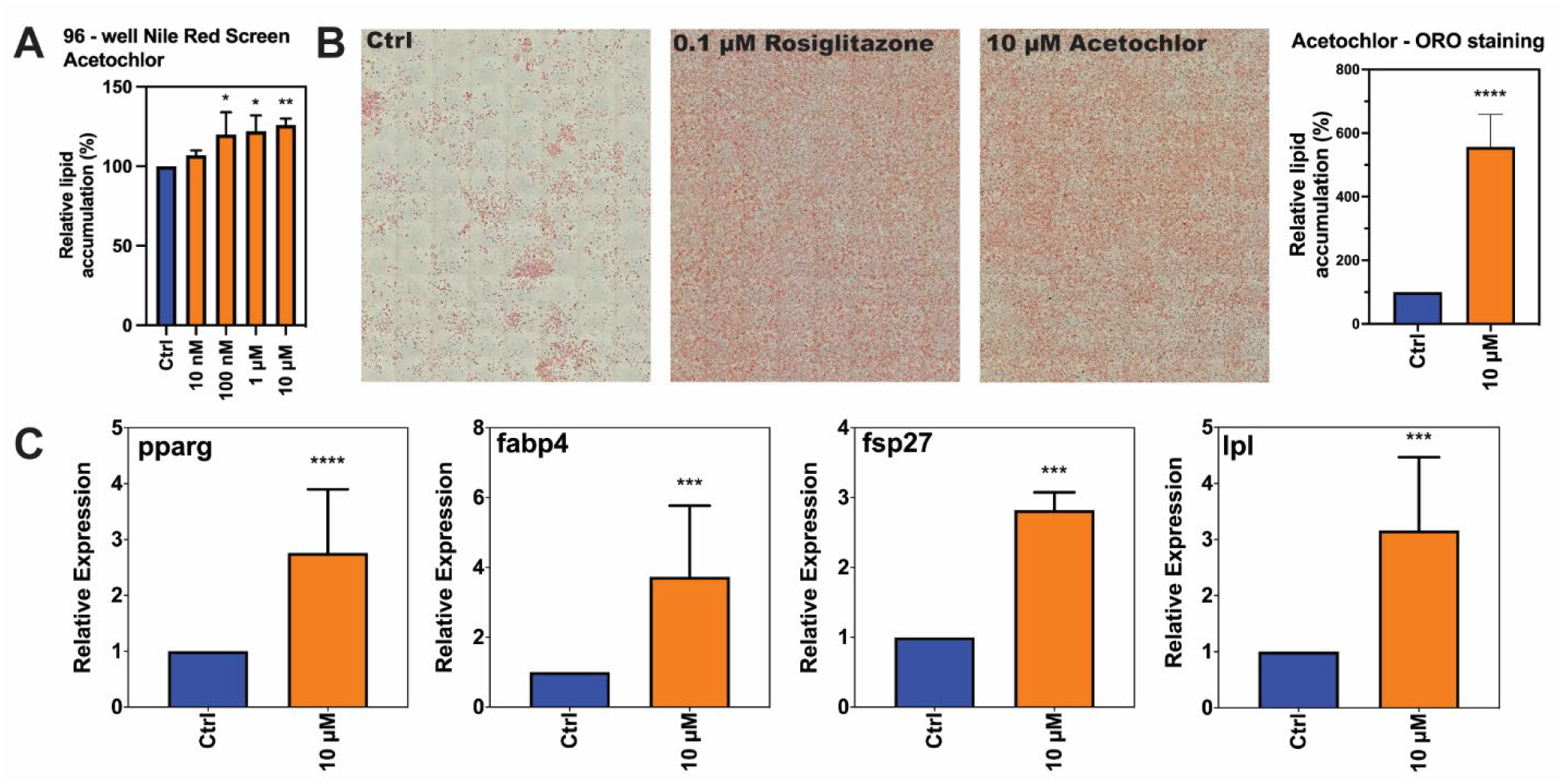
3T3-L1 adipocyte differentiation assay and quantitative PCR. **A)** 3T3L1 cells were cultured for full differentiation in 96 well plate ± acetochlor. Intracellular lipid droplets were stained with AdipoRed and DNA content stained with NucBlue. The lipid contents were normalized to DNA in every well. **B)** Differentiated cells in 6-well plates were stained by Oil-Red-O (ORO). Tile scans at 10X magnification were acquired for each sample and lipid droplets (red stain) were quantified by using ImageJ. **C)** Expression of adipocyte differentiation markers. The expression of adipocyte differentiation markers *Lpl, Fabp4, Pparg*, and *Fsb27* were analyzed by qPCR 8 days post adipogenic induction. Gene expression is presented relative to the value in vehicle control cells. For both **A** and **B**, 0.1 µM of rosiglitazone was used as positive control. The data were normalized to rosiglitazone response and used as input for statistical analysis. One-way ANOVA and Dunnett’s multiple comparisons tests were used to calculate p values. Error bars shown are ± standard deviation of three independent trials. *p < 0.05, **p < 0.01, ***p < 0.001, ****p < 0.0001 for each treatment versus untreated control.

Next, we evaluated the expression of key genes involved in the 3T3-L1 differentiation process by qPCR. Cells treated with 10 µM acetochlor exhibited a significant increase in the expression of *Pparg* (a differentiation marker), *Fabp4* (a preadipocyte marker), *Lpl* (a terminal differentiation marker), and *Fsp27/Cidec* (a lipid droplet marker) [39, 40], indicating that acetochlor promotes differentiation along several steps of the process (**Fig. 1C**). Cells treated with 10 µM of metolachlor had an activation of the late events of adipogenesis, as only the terminal differentiation markers *Lpl* and *Fsp27* were statistically significantly upregulated (**Fig. S3C**, Supporting Information). The induction of *Fsp27* expression is an indicator of strong PPARγ activation [41]; PPARγ being the master regulator of adipogenesis. The other chemicals tested did not induce adipogenesis (**Fig. S4**, Supporting Information). We conclude that acetochlor, and metolachlor are adipogenic chemicals *in vitro*. Since acetochlor and metolachlor both are acetanilide herbicides, we decided to focus on acetochlor only for the follow-up *in vivo* studies to explore acetochlor’s mechanisms of action.

A previous study using 3T3-L1 cells failed to find an effect on adipogenesis when the cells were exposed to 46.7 µM acetochlor [42]. We propose that the lack of observed adipogenicity was caused by general toxicity at this concentration. In our hands, 3T3-L1 cytotoxicity was observed when the cells were treated with 20 µM or higher concentrations of acetochlor or metolachlor (data not shown).

### Acetochlor exposure induces morphological malformations in zebrafish larvae

To investigate obesogenic capacity of acetochlor *in vivo* we turned to the zebrafish model. First, we determined the concentration-response curves for survival and morphological abnormalities resulting from acetochlor exposure.

Acetochlor exposure caused lethality in a concentration-dependent manner with an LC_50_ at 12 µM (**Fig. S5**, Supporting Information). In addition, we observed that it induced general gross malformations in the larvae, such as pericardial edema, curved body axis, and malformed yolk. The effect concentration at which 50% of the surviving larvae had morphological abnormalities (EC_50_) was determined to be 7.8 µM (**Fig. S5**, Supporting Information). The teratogenic index (TI) was calculated as the ratio of LC_50_/EC_50_ to 1.54. According to published criteria used to assess teratogenicity, a chemical with a TI above 1.2 is considered teratogenic [43, 44]. Our lethality and morphology results largely agree with previous studies on acetochlor exposed zebrafish, both regarding the type of malformations observed and the micromolar range of concentrations required to cause malformations and mortality [45, 46].

### Acetochlor exposure induces lipid accumulation in zebrafish larvae

We next set out to determine whether exposure to acetochlor impacts lipid levels *in vivo* in zebrafish larvae. In this study, zebrafish larvae were repeatedly exposed to increasing concentrations of acetochlor under a high cholesterol diet feeding paradigm, schematically shown in **Fig. S2B**, Supporting Information. Several chemicals have previously been found to act as obesogens using this assay, including TBBPA and TCBPA [24], tris(1-chloro-2-propyl)phosphate (TCPP) [47]], perfluorooctanesulfonic acid (PFOS) [48], or a mixture of endocrine disruptors [49].

Acetochlor exposure resulted in increased lipid accumulation in the larvae, seen in the trunk, head and particularly in the blood vessels, compared to unexposed controls (0.005% HFD only) (**Fig. 2A**). Acetochlor exposure elicited a clear dose response, with an increase in accumulated lipids from 1 to 10 nM (**Fig. 2B**). We observed a decrease in ORO staining at higher concentrations of acetochlor (100 nM and 1 µM), likely caused by toxicity and subsequent reduced larval food consumption [50]. Thus, repeated exposure to acetochlor combined with a high cholesterol diet induced lipid accumulation *in vivo*.

**Figure 2.**
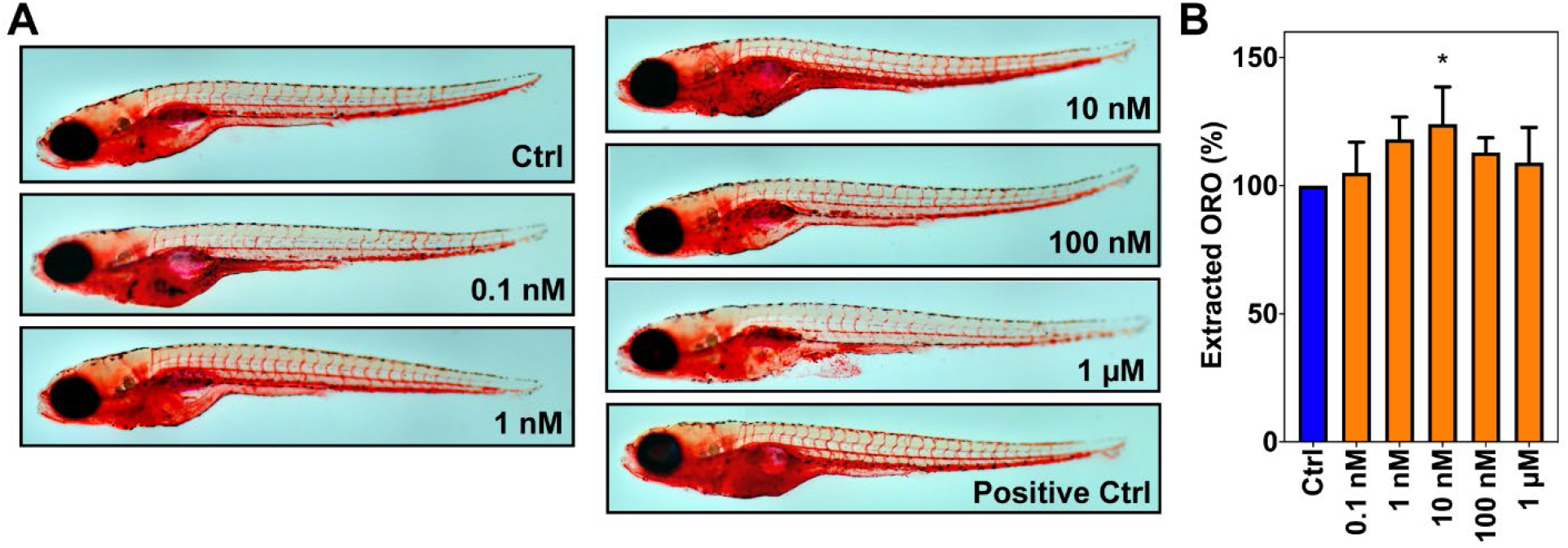
Oil Red O staining of zebrafish larvae. **A)** Representative images of acetochlor treated Oil Red O-stained larvae taken at 5X magnification. The larvae were exposed from 0.1 nM to 1 µM acetochlor and fed with 0.005% HFD. Control is 0.005% HFD only. The positive control is 0.1% HFD without exposure. **B)** Quantification of ORO staining in zebrafish larvae. The extracted ORO was measured at 495 nM and data were presented relative to the value in 0.005% HFD. Normalized data to 0.1% HFD positive control response was used as input for statistical analysis. Unpaired two-tailed Student t-test was used to calculate p values. Error bars shown are ± standard deviation of three independent trials. *p < 0.05 for sample versus untreated control.

### Acetochlor exposure alters the expression of adipogenic-signature genes in zebrafish larvae

To investigate modes of action related to metabolic endpoints altered by acetochlor we performed mRNA sequencing of exposed whole zebrafish larvae. Forty-eight hpf zebrafish embryos were exposed to 7.8 µM (EC_50_) acetochlor for 24 h, followed by mRNA isolation and sequencing. Unsupervised hierarchical clustering by Euclidian distance demonstrated a clear separation between treated and untreated replicates (**Fig. S6A**, Supporting Information). The analysis showed that 1211 genes were differentially expressed: 754 genes were upregulated, and 457 genes were downregulated (**Fig. S6B**, Supporting Information).

We conducted upstream analysis to predict the top-10 transcription factors (TFs) that were likely responsible for the observed changes in gene expression (**Fig. 3A**). Several of the predicted enriched transcription factors are known to play a role in regulating different aspects of cellular metabolism, including *tp53* [51], *nrf2* **(Fig. 3B and C)** [52], *ppara* [53, 54] and the transcription coactivator *ppargc1a/PGC-1alpha* [55]. PPARα is known to play a crucial role in regulating lipid homeostasis [56], and PGC-1alpha is involved in mitochondrial biogenesis and oxidative phosphorylation, gluconeogenesis, and fatty acid synthesis [57]. NFE2L2/NRF2 is a transcription factor that binds to antioxidant response elements (AREs) to induce the expression of several antioxidant and detoxification genes [58]. Thus, NRF2 is important to maintain redox homeostasis and to mitigate the adverse effects of oxidative stress, such as those resulting from lipid peroxidation and ferroptosis [59]. Furthermore, in response to oxidative stress, NRF2 promotes lipid accumulation in adipocytes by stimulating lipogenesis and inhibiting lipolysis [60].

**Figure 3.**
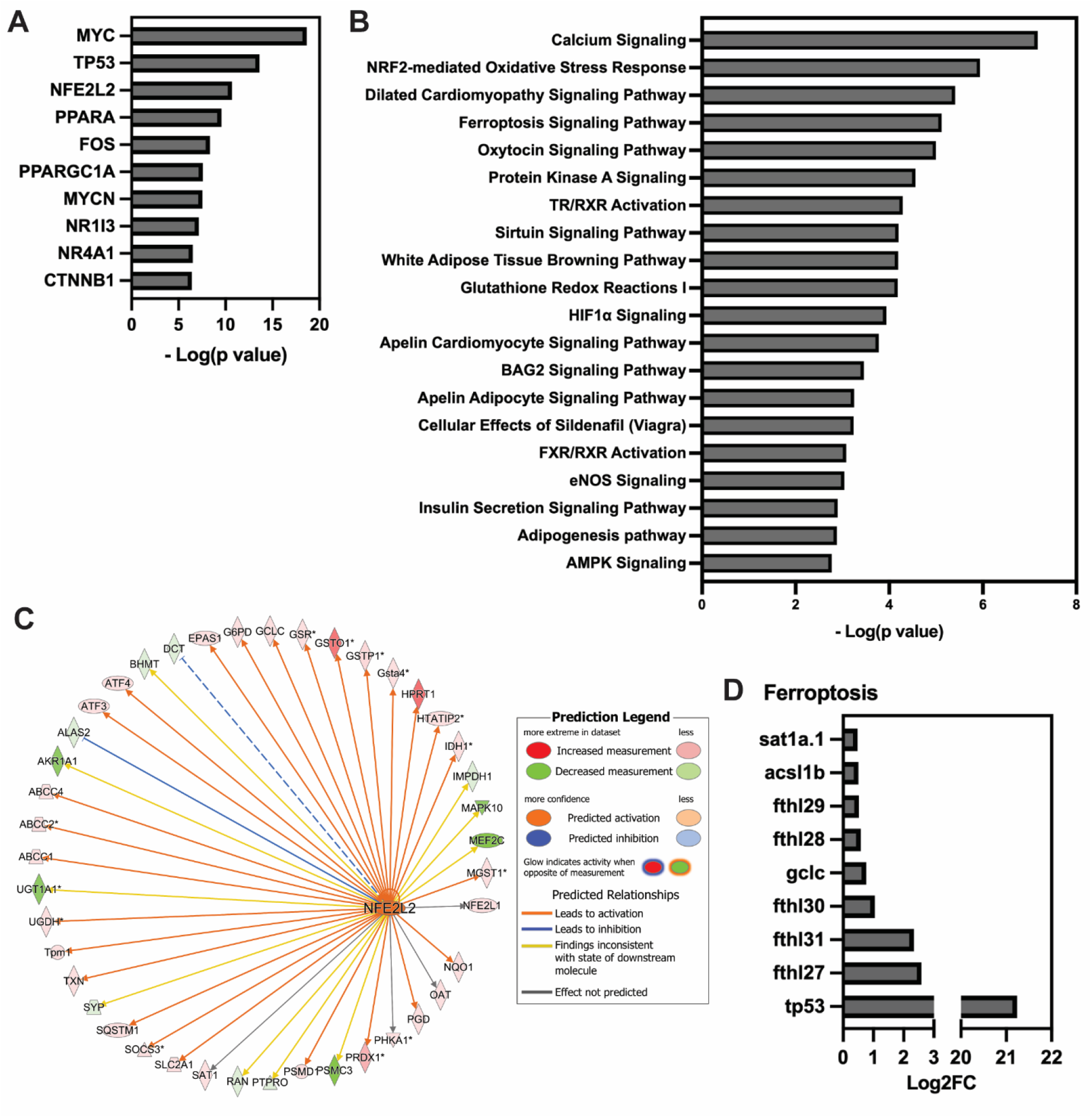
Pathway analysis of acetochlor treated zebrafish larvae. At 48 hpf, zebrafish larvae were treated with 7.8 µM acetochlor and RNA were collected at 72 hpf. **A)** Enriched transcription factors as potential regulators of DEGs predicted by IPA. **B)** Top 20 canonical pathways by analysis using IPA. **C)** Predicted activation of genes downstream of the oxidative stress factor NRF2/NFE2-L2. **D)** Enrichment in ferroptosis regulated genes, including several ferritin heavy chain-like (fthl) genes.

We next examined major biological functions and cellular processes affected by acetochlor exposure. IPA analysis predicted that processes related to lipid, glucose, and monosaccharide metabolisms, as well as adipogenesis, such as CEBPA[61] and CREB3L3-regulated[62] pathways were upregulated (**Fig. 3B and Fig. S6C**, Supporting Information). The top canonical enriched pathways included calcium signaling and, again, NRF2-related responses. Induction of calcium signaling is known to be associated with endoplasmic reticulum stress [63, 64] and mitochondrial dysfunction [63, 65]. IPA also predicted an enrichment in ferroptosis regulated genes (**Fig. 3D**). Ferroptosis is an iron-dependent, oxidative stress related cell death caused by membrane lipid peroxides [66]. Elevated mitochondrial oxidative stress, reactive oxygen species (ROS), and lipid peroxides are associated with an obesity phenotype [67, 68]. The thyroid hormone receptor/retinoid X receptor signaling pathway (TR/RXR) was also predicted to be altered by the acetochlor exposure (**Fig. 3C**), which is in accordance with previously published reports in fish and frogs [69-71]. Further examination of the NRF2, PPARGC1A, and PPARα regulated genes (**Fig. S5D**, Supporting Information) identified downstream genes known to be involved in lipid metabolic pathways, such as *hmgcs1, fabp2, cyp27a1* and *cyp8b1* (**Fig. S5E**, Supporting Information).

In summary, the transcriptomic results suggest that acetochlor exposure induces ROS production, because *Nrf2* driven gene expression was strongly activated. It further suggested that exposure altered major pathways involved in metabolic homeostasis and adipose tissue development and/or function, including PGC-1alpha and thyroid hormone signaling. Thus, the activities of acetochlor are comparable to those of obesogens [72-74]. Furthermore, our transcriptomic analysis suggested that ferroptosis was induced. The acetochlor effect seen here is in line with a reported impaired fatty acid oxidation by acetochlor in mice [75]. Additionally, long-term exposure to acetochlor promotes DNA damage caused by oxidative stress in Bufo raddei tadpoles [76]. Finally, elevated levels of ROS in mesenchymal stem cells are known to favor adipogenesis over osteogenesis [77].

### Acetochlor exposure results in lipid peroxidation and inhibition of glutathione peroxidase

To explore the changes in the lipidome following exposure of zebrafish larvae to acetochlor, we analyzed the abundance of 626 annotated lipids. The lipid species identification is based on the fact that zebrafish shares remarkable conservation in lipid biochemistry with humans [78]. Additionally, the zebrafish embryo is lecithotrophic, relying exclusively on its yolk repository of proteins and lipids to support early embryonic growth [79]. This lecithotrophic state prior to the transition to free-feeding animal reduces experimental variabilities that arise from exogenous feeding [80]. These features make larval zebrafish a well-suited organism to assess chemical influences on vertebrate lipid metabolism.

Acetochlor exposure induced a distinct, although not drastic, deviation in lipid content compared to control as depicted in the PLS-DA plot (**Fig. S7A**, Supporting Information). This modest effect might be explained by our short exposure time of 24 h; a larger effect on the lipid profile might require longer exposure times. Nevertheless, we found significant increase in ceramide levels (**Fig. 4A and B**, and **Fig. S7B**, Supporting Information), possibly resulting from increased ROS and/or generation of oxidized phospholipid [81, 82]. High levels of ceramides are found in individuals with obesity and dyslipidemia, resulting in poor metabolic outcomes, such as insulin resistance and triglyceride synthesis [83]. We also found an increase in acylcarnitines (ACar) (**Fig. 4A and B**, and **Fig. S7B**, Supporting Information) indicating dysfunction in β-oxidation and catabolism of long-chain FA [84]. Finally, the lipidomics data also revealed an elevation of oxidized phosphatidylethanolamine (OxPE) levels (**Fig. 4A and B**, and **Fig. S7B**, Supporting Information).

**Figure 4.**
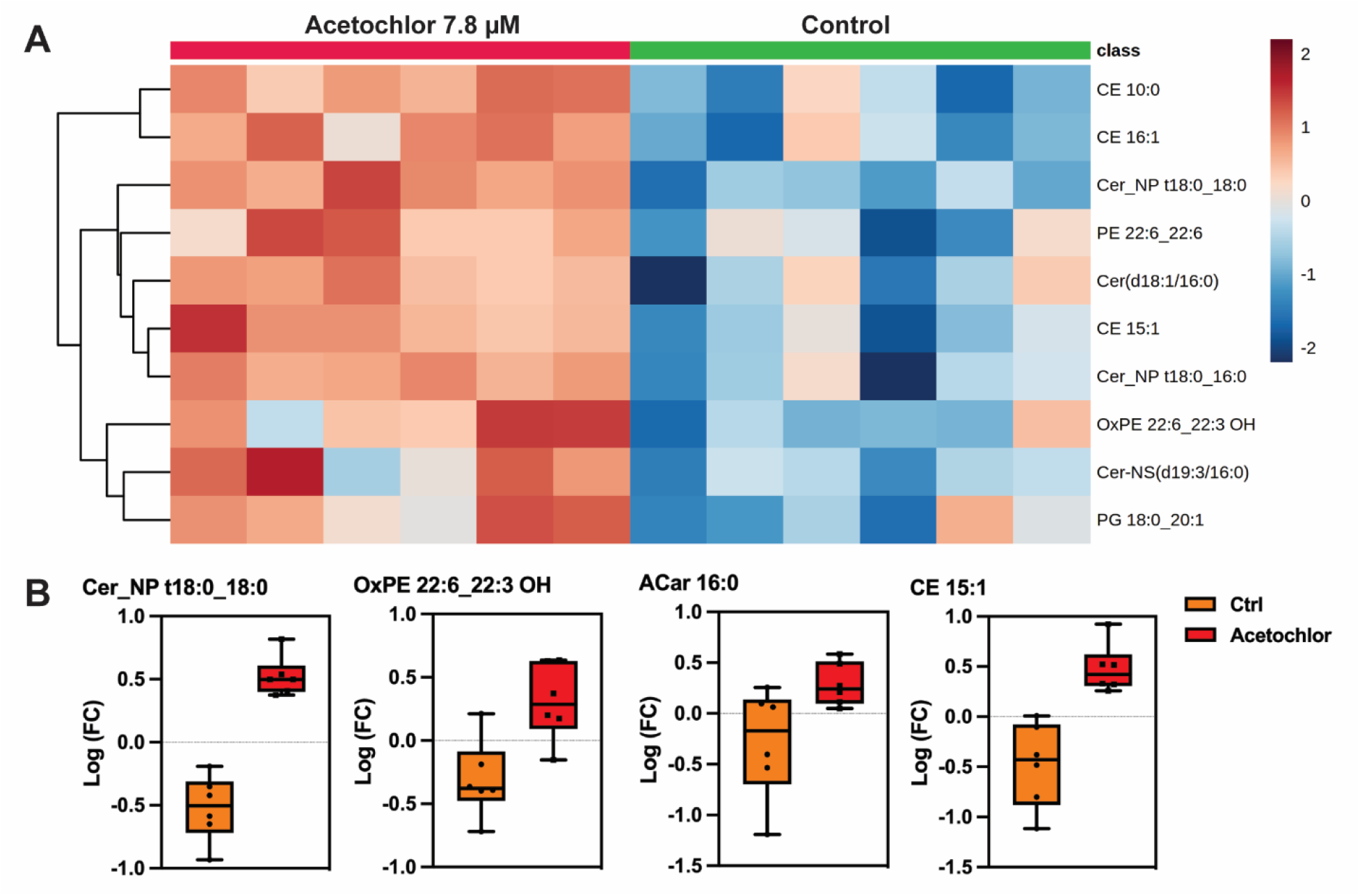
Altered lipids in zebrafish larvae treated with acetochlor revealed by lipidomics. At 48 hpf zebrafish larvae were treated with 7.8 µM acetochlor and snap frozen at 72 hpf for lipidomic analysis. **A)** Heatmap of lipids for 6 replicates of control and treatment based on normalized mass spectrometry (MS) intensities of the top 10 lipid classes as determined by one-way ANOVA and post-hoc analysis. **B)** Log fold change (FC) of the normalized MS intensities for lipids of the classes of ceramides (Cer), oxidized phosphatidylethanolamines (OxPE), acylcarnitines (ACar), and cholesteryl esters (CE).

The transcriptomic and lipidomic results combined inferred an increased Nrf2 activity in response to ROS, and induced ferroptosis and lipid peroxidation after acetochlor exposure. This combination of effects suggested that acetochlor interferes with essential regulatory proteins of ferroptosis. Most ferroptosis inducers function by inhibiting glutathione peroxidase 4 (GPX4) or ferroptosis suppressor protein 1 (FSP1) [85]. The selenoenzyme GPX4 reduces membrane lipid hydroperoxides into non-toxic lipid alcohols using glutathione as a substrate. GPX4 knockout mice are embryonic lethal, demonstrating that its activity is necessary for life [86]. Interestingly, we found that acetochlor has a structural similarity with the known human GPX4 inhibitors RSL3 and ML162 (**Fig. 5A**). The shared chloroacetamide group is an essential chemical moiety that covalently binds to GPX4, and directly inhibits its catalytic activity in an irreversible manner [85].

**Figure 5.**
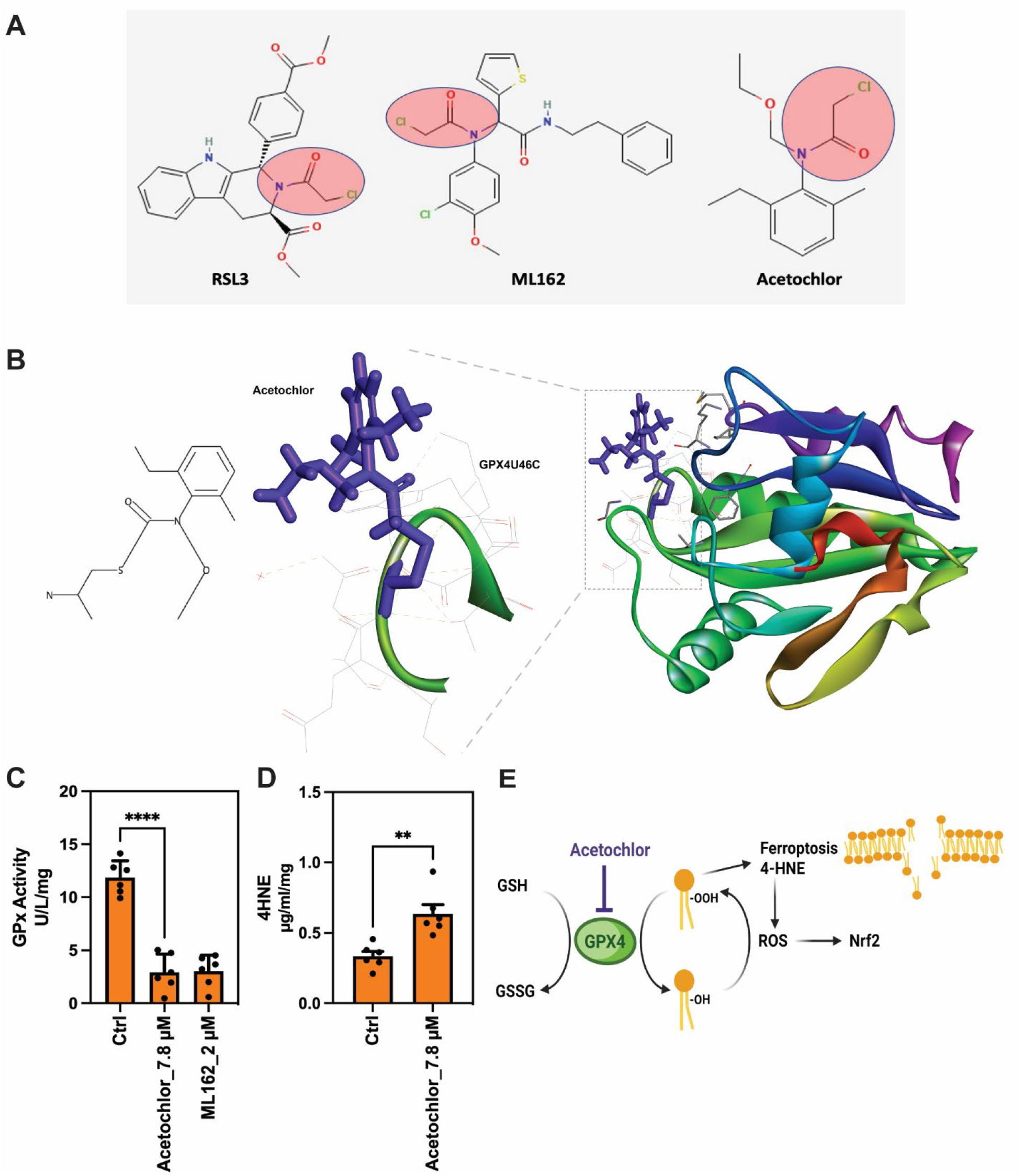
Acetochlor inhibits glutathione peroxidase 4 activity and induces lipid peroxidation in zebrafish larvae. **A)** Structural similarity of acetochlor and the known GPX4 inhibitors RSL3 and ML162. **B)** Simulation of the covalent interaction between GPX4U46C (green) [PDB:2OBI] and acetochlor (purple) using the Forecaster Suite **C)** Acetochlor exposure at 7.8 µM inhibits GPX activity as efficiently as ML162 at 2 µM. **D)** Acetochlor exposure at 7.8 µM induces 4-HNE accumulation. **E)** The proposed models for the effect of acetochlor mediated disruption lipid homeostasis (created with BioRender.com). Unpaired two-tailed Student t-test was used to calculate p values in **C** and **D**. Error bars shown are ± standard deviation of three independent trials. **p < 0.01 and ****p < 0.0001 for sample versus untreated control.

To predict an acetochlor - GPX4 interaction, a covalent docking simulation study was performed using the small molecule FITTED docking approach [34-36]. As the software only recognizes standard amino acids, selenocysteine residue 46 (Sec/U) in GPX4 was replaced with cysteine in the active site. Interestingly, the molecular docking algorithm predicted a covalent bond formation between acetochlor and this cysteine, forming a covalent protein-inhibitor complex (**Fig. 5B**). Naturally, selenocysteine exerts a higher nucleophilic activity compared to cysteine [87], suggesting that a strong selenocysteine – acetochlor covalent interaction would occur. We also note that the selenocysteine in GPX4 and several amino acids of its active site are conserved between human, mouse, and zebrafish GPX4s (**Fig. S7A**, Supporting Information). Moreover, the selenocysteine in the active site is also conserved between different glutathione peroxidases (**Fig. S7B**, Supporting Information).

To validate the proposed inhibitory function of acetochlor on Gpx4 experimentally, we exposed zebrafish larvae to acetochlor for 24 h followed by analysis of Gpx4 activity. Acetochlor (7.8 µM) exposure led to an 80% reduction in Gpx4 activity and had as strong an inhibitory activity as 2 µM ML162, indicating an actual inhibitory acetochlor selenocysteine – Gpx4 interaction (**Fig. 5C**). Next, we investigated the effect of acetochlor exposure on 4-hydroxynonenal (4-HNE) levels, a major biomarker for lipid peroxidation. 4-HNE modulates multiple signaling pathways, including induction of Nrf2, AP-1 and PPAR activities [88-90] or inhibition of SIRT [91]. Moreover, 4-HNE itself induces mitochondrial oxidative stress is several tissues [92]. In this experiment we extended the acetochlor exposure time to four days to allow for a stronger induction of lipid peroxidation. We found a significant increase in 4-HNE levels in the acetochlor treated larvae compared to untreated controls (**Fig. 5D**).

Based on these results, we propose a model for how acetochlor disrupts lipid homeostasis (**Fig. 5E**). By inhibiting Gpx4, acetochlor exposure results in excessive lipid peroxidation and 4-HNE production. Likely depending on the exposure concentration, acetochlor will in this way cause cell death by ferroptosis, or oxidative stress-induced promotion of adipogenic pathways. In the long run, increased lipid peroxidation, 4-HNE levels, and ROS, cause metabolic dysregulation and development of metabolic disorders [92, 93]. We propose that this molecular mechanism is one of the main obesogenic activities of acetochlor, but we do not exclude that it also interacts with other enzymes and pathways. We note that GPX4 in addition to reducing lipid peroxidation, also protects against mitochondrial-derived ROS in spermatocytes and is absolutely required for male fertility [94]. Moreover, loss of GPX4 is implicated in neurodegeneration [95]. We also note that metolachlor, among other acetanilide herbicides, shares the same chemical moiety as acetochlor, RSL3, and ML162, indicating that it may also inhibit Gpx4 activity. Thus, we suggest that inhibition of Gpx4 is a novel molecular initiating event by obesogenic chemicals.

## Abbreviations

(ORO): Oil Red O
(GPX): Glutathione peroxidase
(dpf): Days post fertilization
(hpf): Hours post fertilization
(4-HNE): 4-Hydroxynonenal
(Nrf2): Nuclear factor erythroid 2–related factor 2
(DMSO): dimethylsulfoxide
(PBS): phosphate buffered saline
(IPA): Ingenuity Pathway Analysis
(TI): teratogenic index
(LC): lethal concentration
(EC): effect concentration
(ROS): reactive oxygen species
(ACar): acylcarnitines
(OxPE): oxidized phosphatidylethanolamine

## Acknowledgments

We thank the Rural Obesity Project Development Team at Indiana University for fruitful discussions of the study. We thank Nicolas Moitessier for guidance on the Molecular Forcaster Suite software. Metabolomics analysis was performed by John Allen Mack in the Metabolomics Core at the University of Utah School of Medicine Centers using equipment provided for by University of Utah RIF funds. RNA sequencing was performed at the Center for Genomics and Bioinformatics at Indiana University Bloomington.

## Supporting Information

Supporting Materials and Methods

Table S1. Primer sequences for qPCR.

Figure S1. Molecular structures of investigated pesticides.

Figure S2. Schematic depiction of the experimental procedures.

Figure S3. 3T3-L1 adipocyte differentiation assay and quantitative PCR after metolachlor exposure.

Figure S4. 3T3-L1 adipocyte differentiation assay and quantitative PCR after pesticide exposure. Figure S5. Lethality and teratogenic effects in zebrafish larvae by exposure to acetochlor.

Figure S6. RNA sequencing and pathway analysis of acetochlor treated zebrafish larvae.

Figure S7. Lipid signatures in zebrafish larvae treated with acetochlor revealed by lipidomics.

Figure S8. The selenocysteine of GPX4 is conserved through species and GPXs.

References for Supporting Information

