## Supporting Information for "The herbicide acetochlor causes lipid peroxidation by inhibition of glutathione peroxidase 4"

### Materials and Methods

**Chemicals**

All chemicals were of PESTANAL® analytical or HPLC grade and were purchased from Millipore Sigma (St. Louis, MO) as follows: 2-chloro-N-ethoxymethyl-N-(2′-ethyl-6′-methylphenyl) acetamide; acetochlor (CAS# 34256-82-1), 2-chloro-4-ethylamino-6-isopropylamino-1,3,5-triazine; atrazine (CAS# 1912-24-9), 3,6-dichloro-2-methoxybenzoic acid, 3,6-dichloro-o-anisic acid; dicamba (CAS# 1918-00-9), N-(2,6-difluorophenyl)-5-methyl-[1,2,4]triazolo[1,5-a]pyrimidine-2-sulfonamide, 2′,6′-difluoro-5-methyl-[1,2,4]triazolo[1,5-a]pyrimidine-2-sulfonanilide; flumetsulam (CAS# 98967-40-9), N-(phosphonomethyl)glycine; glyphosate (CAS# 1071-83-6), 2-chloro-N-(2-ethyl-6-methylphenyl)-N-[(1S)-2-methoxy-1-methylethyl]acetamide; metolachlor, predominantly S-metolachlor (CAS# 87392-12-9), 5-[[4-[2-(methyl-2-pyridinylamino)ethoxy]phenyl]methyl]-2,4-thiazolidinedione; rosiglitazone (CAS# 155141-29-0) and α-[(2-chloroacetyl)(3-chloro-4-methoxyphenyl)amino]-N-(2-phenylethyl)-2-thiopheneacetamide, 2-chloro-N-(3-chloro-4-methoxyphenyl)-N-(2-oxo-2-(phenethylamino)-1-(thiophen-2-yl)ethyl)acetamide; ML162 (CAS# 1035072-16-2). The stock solutions were prepared in dimethylsulfoxide (DMSO) (CAS# 67-68-5) except glyphosate, which was prepared in phosphate buffered saline; PBS (pH 7.4). To avoid freeze-thaw cycles, stocks were stored as aliquots at -20 $^{0}$C until use. The final dilutions of these stock solutions to be used in the cell cultures or zebrafish treatments always resulted in a DMSO concentration below 0.1%.

**3T3-L1 cell differentiation**

Cells were harvested in 1X Trypsin-EDTA (Thermo Fisher, Waltham, MA) and seeded in 96 well plates at $2x10^3$ cells/well or in 6 well plates at $8x10^4$ cells/well in preadipocyte expansion media and cultured until 100% confluency (48-72 h). Pre-adipocyte expansion media was renewed, and cells were maintained as a confluent culture for 48 h to initiate growth arrest and clonal expansion. Media was then replaced with differentiation media consisting of DMEM-HG supplemented with 10% Fetal Bovine Serum; FBS (GE HealthCare, Chicago, IL), 1% anti-anti, 0.5 µM dexamethasone (Millipore Sigma), 0.25 mM isobutylmethylxanthine; IBMX (Sigma; I7018), and 5 µg/ml insulin (Millipore Sigma). The test chemicals or vehicle controls were included with the differentiation media. After 48 h, the media was replaced with adipocyte maintenance media consisting of DMEM-HG, 10% FBS, 1% anti-anti, and 5 µg/ml insulin in the presence of test chemicals or vehicle and this media was refreshed every 48 h.

***In vitro* 3T3-L1 Nile red and NucBlue staining**

The 3T3-L1 cells were seeded into 96-well, clear bottom black cell culture microplates (Greiner Bio-One, Monroe, NC) and differentiated as described above with the indicated treatments for 8 days. The media was gently removed, and the cells were rinsed 3-4 times with PBS. Nile red (AdipoRed; Lonza Bioscience, Walkersville, MD) was used to quantify intracellular lipids and NucBlue live ready probe (Thermo Fisher) was used to quantify DNA content. The dyes were prepared by mixing 20 drops of NucBlue into 18.5 ml of PBS and 500 µl AdipoRed, and 200 µl of the dye mixture was used per well. After incubation of the microplates for 40 minutes in darkness at room temperature, the fluorescence was measured by a microplate reader (BioTek Synergy H1; Agilent Technologies, Santa Clara, CA) with excitation/emission 485/572 for AdipoRed and 360/460 for NucBlue. Finally, Nile red fluorescence was normalized by cell number (DNA content) to accurately define the adipogenic activity of the assayed compounds.

**Zebrafish husbandry**

All animal work was approved by the Indiana University, Bloomington Institutional Animal Care and Use Committee. Adult zebrafish (*Danio rerio*) were maintained in tanks in an Aquaneering system (Aquaneering USA Inc., San Diego, CA) supplied with circulating reverse osmosis water at 28 ± 0.5 °C in a 14/10 h light/dark cycle (lights on 8 AM; lights off 10 PM). The fish were fed commercial flake food (TetraMin; Tetra, The Woodlands, TX) in the morning and baby brine shrimp (Brine Shrimp Direct, Ogden, UT) in the afternoon. Adult fish were set up for breeding in the late afternoon and the embryos were collected in the following morning and maintained in embryo media (E3; 5 mM NaCl, 0.17 mM KCl, 0.33 mM $CaCl_2$, 0.33 mM $MgSO_4$ adjusted to pH 7.4) at 28.5 ± 0.5 °C.

***In vivo* exposure and lipid accumulation assay**

Wild type NHGRI-1 fish (Zebrafish International Resource Center (ZIRC) at the University of Oregon, OR) were placed in breeding tanks overnight. The next day, the embryos were collected in 10 cm plates and selected for the exposure. From day 1 to 5, the fish were treated with 1X phenylthiourea (PTU) (Millipore Sigma) in E3. At day 3, we started acetochlor exposure at 0.1 nM, 1 nM, 10 nM, 100 nM and 1 µM concentrations, and 0.01% DMSO was used as vehicle control.

At day 6, the larvae were moved to the feeding tanks that contain the chemical or vehicle control (DMSO) in 350 ml E3 mixed with 0.005% (w/v) high fat diet (HFD; chicken egg yolk powder (Magic Flavors; Seattle, WA). The larvae were fed for 6 h, then the exposure medium was replaced with fresh E3 with replenished chemical treatment. We repeated the feeding and treatment paradigm until day 10. After day 10, the larvae did not receive any food for 24 h allowing them to clear their gut [1] . The larvae were euthanized at day 11 and fixed in 4% PFA at $4^{o}$C for at least 24 hr. ORO larval staining was performed as described before [2] Briefly, the larvae were rinsed and permeabilized in 60% isopropanol for 1 h at room temperature, stained with ORO working solution (as described above) for 75 minutes. The stained larvae were rinsed twice with 0.1% PBST (0.1% Tween-20 in PBS), washed with a series of glycerol solutions; 25% glycerol (made in 0.1% PBST), 50% glycerol and 75% glycerol, each for 15 minutes at room temperature. Embryos were stored in 75% glycerol at $4^{o}$C. The ORO staining was performed three times with at least 20 larvae.

**Targeted lipidomics**

Extraction of lipids from 72 hpf zebrafish larvae was carried out using a biphasic solvent system of cold methanol, methyl tert-butyl ether (MTBE), and PBS/water based on Matyash *et al*. with some modifications [3]. Lipid extracts were separated on an Acquity UPLC CSH C18 column (2.1 x 100 mm; 1.7 µm) coupled to an Acquity UPLC CSH C18 VanGuard precolumn (5 × 2.1 mm; 1.7 µm) (Waters; 186003980) maintained at 65°C connected to an Agilent HiP 1290 Sampler, Agilent 1290 Infinity pump, and Agilent 6545 Accurate Mass Q-TOF dual AJS-ESI mass spectrometer (Agilent Technologies). Samples were analyzed in a randomized order in both positive and negative ionization modes in separate experiments acquiring with the scan range m/z 100 – 1700. Six replicates were analyzed for both control and exposed larvae.

For data processing, Agilent MassHunter (MH) Workstation and software packages MH Qualitative and MH Quantitative were used. For lipid annotation, accurate mass and MS/MS matching was used with the Agilent Lipid Annotator library and LipidMatch [4]. Results from the positive and negative ionization modes from Lipid Annotator were merged based on the class of lipid identified. Multivariate analysis was performed using MetaboAnalyst [5]. Statistical models were created for the normalized data after normalizing to sum, logarithmic transformation (base 10) and Pareto scaling. A p-value of $< 0.05$ and a FC $\geq 1.2$ were used for heatmaps or log fold change analysis. For lipid ontology and enrichment analysis, LION web was used to assess the changes in the sets of lipids that share a certain property [6].

**Table S1. Primer sequences for qPCR**

| Gene [mouse nomenclature] | Sequence of the primer 5'-3' | |
|---|---|---|
| | **Forward** | **Reverse** |
| **lpl** | CATCGAGAGGATCCGAGTGAA | TGCTGAGTCCTTTCCCTTCTG |
| **fabp4** | GATGAAATCACCGCAGACGAC | ATTCCACCACCAGCTTGTCAC |
| **fsp27** | CTGGAGGAAGATGGCACAAT | GGGCCACATCGATCTTCTTA |
| **pparg** | CATAAAGTCCTTCCCGCTGA | GAAACTGGCACCCTTGAAAA |
| **arhgdia** | AAGGACGATGAAAGCCTCCG | GGTCAGTCGAGTCACAATGACA |
| **actb** | GGCTGTATTCCCCTCCATCG | CCAGTTGGTAACAATGCCATGT |

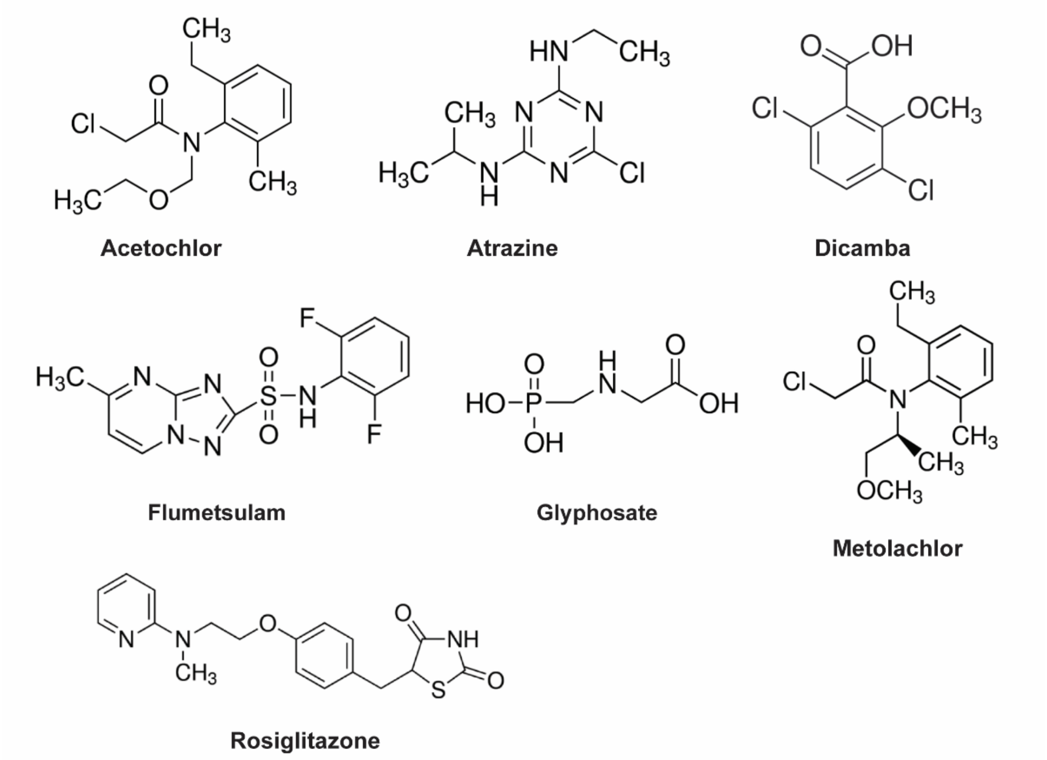


**Figure S1. Molecular structures of investigated pesticides.** Acetochlor, atrazine, dicamba, flumetsulam, glyphosate, and metolachlor are herbicides. Rosiglitazone is an PPARγ agonist. The commercial names are: atrazine (sold as AAtrex), metolachlor (Dual), acetochlor (Harness, Topnotch), glyphosate (Ranger, Rattler, Rodeo, Roundup), dicamba (Banvel), and flumetsulam (Broadstrike). Structures are from Sigma Aldrich's structure search (https://www.sigmaaldrich.com/US/en/structure-search).

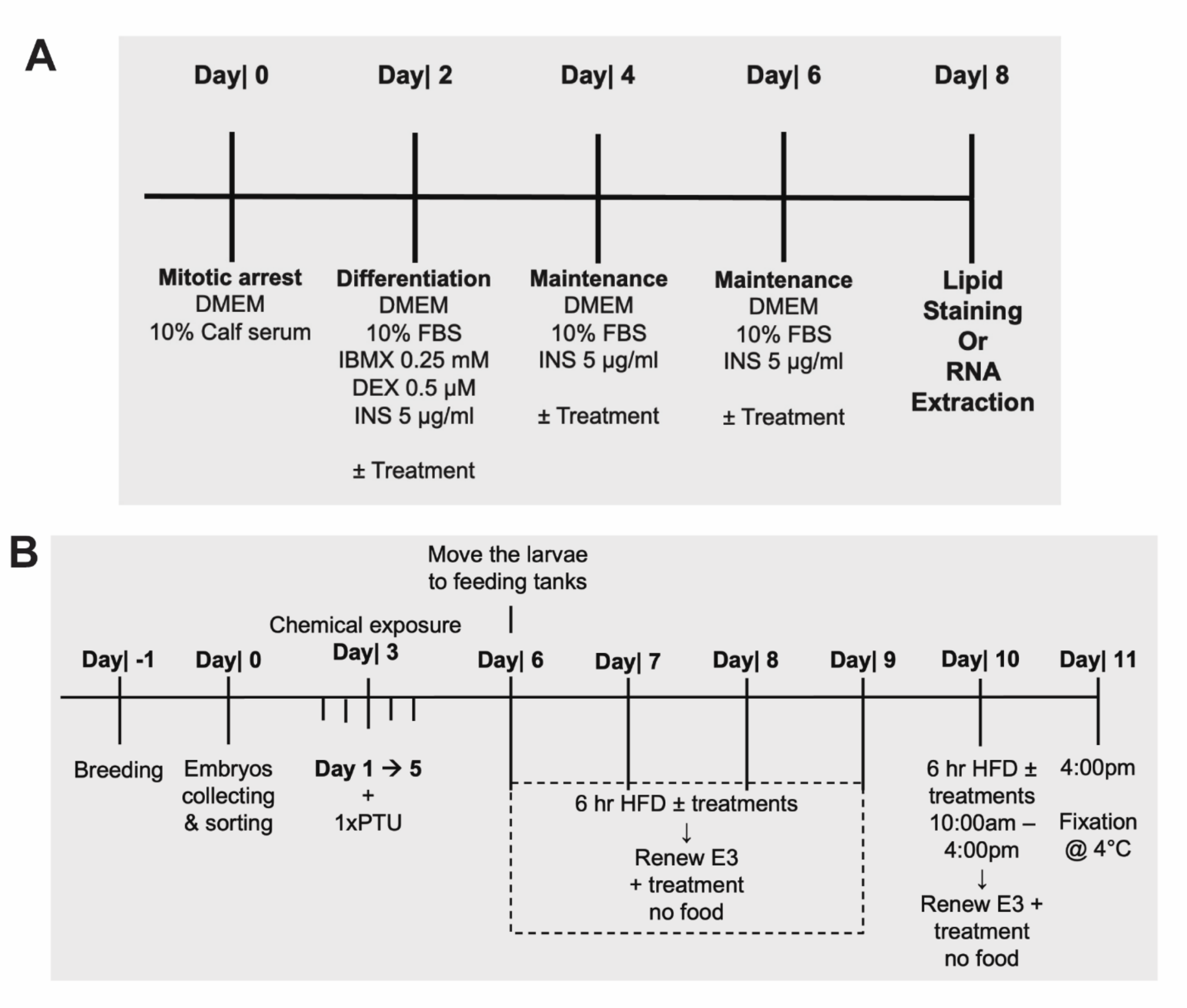


**Figure S2. Schematic depiction of the experimental procedures. A)** 3T3-L1 cell differentiation to adipocytes, and **B)** lipid accumulation assay in zebrafish larvae.

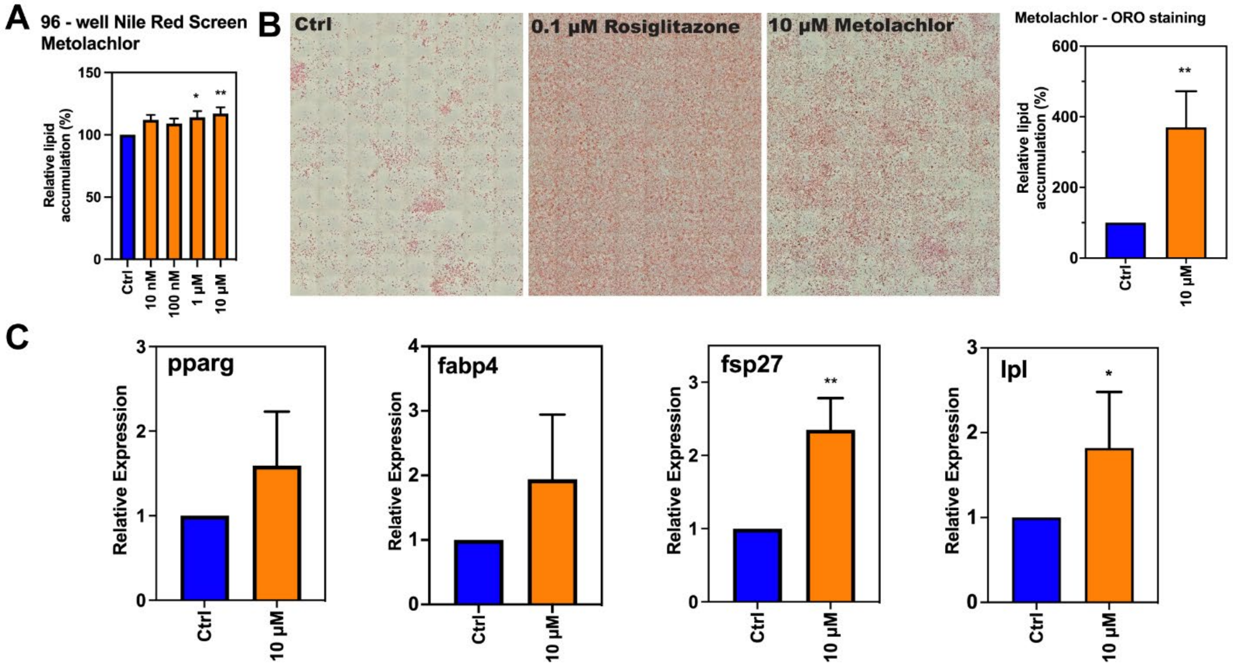


**Figure S3. 3T3-L1 adipocyte differentiation assay and quantitative PCR after metolachlor exposure. A)** 3T3L1 cells were cultured for full differentiation in 96 well plate ± ametolachlor. Intracellular lipid droplets were stained with AdipoRed and DNA content stained with NucBlue. The lipid contents were normalized to DNA in every well. **B)** Differentiated cells in 6-well plates were stained by Oil-Red-O (ORO). Tile scans at 10X magnification were acquired for each sample and lipid droplets (red stain) were quantified by using ImageJ. **C)** Expression of adipocyte differentiation markers. The expression of adipocyte differentiation markers *lpl*, *fabp4*, *pparg*, and *fsb27* were analyzed by qPCR 8 days post adipogenic induction. Gene expression is presented relative to the value in vehicle control cells. 0.1 µM of rosiglitazone was used as positive control in all experiments. The data were normalized to rosiglitazone response and used as input for statistical analysis. One-way ANOVA and Dunnett's multiple comparisons tests were used to calculate p values. Error bars shown are ± standard deviation of three independent trials. *p < 0.05, **p < 0.01, for each treatment versus untreated control (Ctrl).

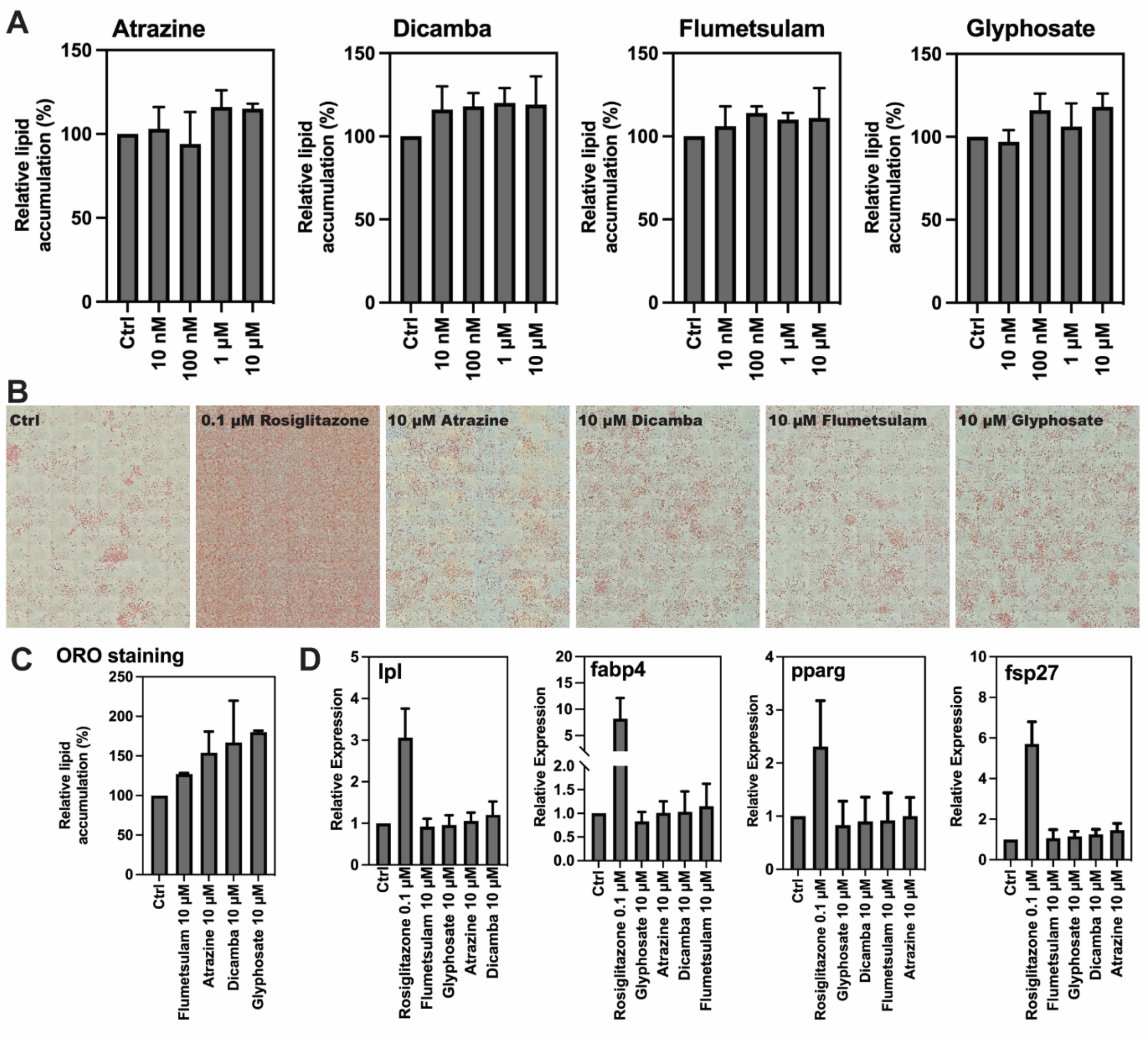


**Figure S4. 3T3-L1 adipocyte differentiation assay and quantitative PCR after pesticide exposure. A)** 3T3L1 cells were cultured for full differentiation in 96 well plate ± pesticides. Intracellular lipid droplets were stained with AdipoRed and DNA content stained with NucBlue. The lipid contents were normalized to DNA in every well. **B)** Differentiated cells in 6-well plates were stained by Oil-Red-O (ORO). Tile scans at 10X magnification were acquired for each sample and lipid droplets (red stain) were quantified by using ImageJ. **C)** Expression of adipocyte differentiation markers. The expression of adipocyte differentiation markers *lpl*, *fabp4*, *pparg*, and *fsb27* were analyzed by qPCR 8 days post adipogenic induction. Gene expression is presented relative to the value in vehicle control cells. In all experiments, 0.1 µM of rosiglitazone was used as positive control. The data were normalized to rosiglitazone response and used as input for statistical analysis. One-way ANOVA and Dunnett's multiple comparisons tests were used to calculate p values. Error bars shown are ± standard deviation of three independent trials. None of the exposures resulted in statistically significant changes compared to control (Ctrl).

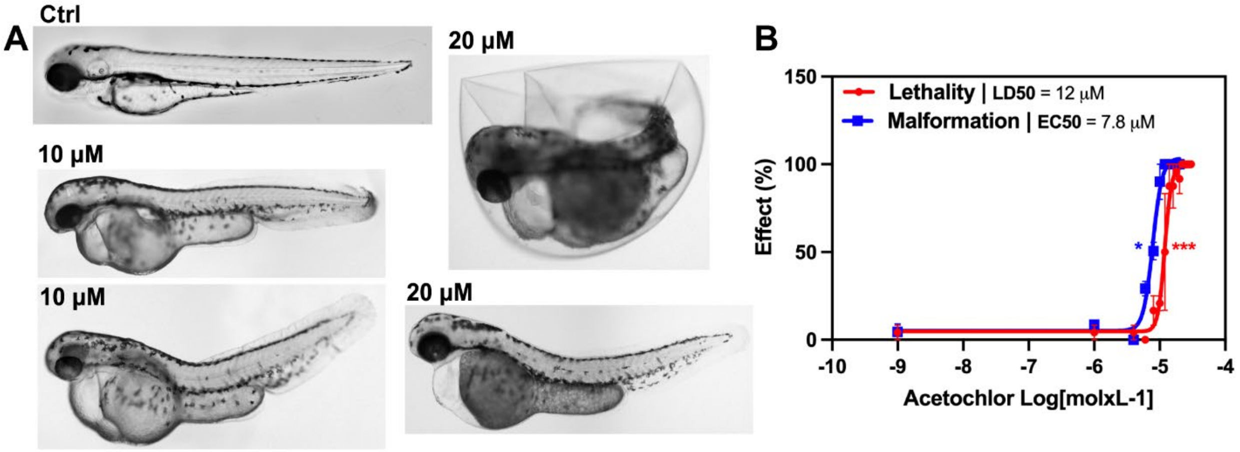


**Figure S5. Lethality and teratogenic effects in zebrafish larvae by exposure to acetochlor. A)** Examples of morphological malformations after acetchlor exposure. **B)** Concentration response curves for malformations and lethality in acetochlor exposed embryos. The zebrafish embryos were arrayed in 96-well plates, treated with increasing concentrations of acetochlor at 6 hpf, and analyzed at 72 hpf. The addition of the chemical at the onset of gastrulation (shield stage = 6 hpf) allows for it to interfere with critical stages of embryonic development and cell fate specification, mimicking mammalian developmental toxicity studies [7]. The lethal concentration 50% (**$LC_{50}$**) was 12 µM and the malformation concentration 50% (**$EC_{50}$**) was 7.8 µM. A teratogenic index (**TI = $LD_{50}/ED_{50}$**) was calculated to be 1.54, indicating teratogenic potential of acetochlor on zebrafish embryos under the experimental conditions used. Non-parametric Kruskal–Wallis and Dunn's multiple comparisons tests were used to calculate p values. *$p < 0.05$, **$p < 0.01$, ***$p < 0.001$ versus untreated control.

Acetochlor has previously been shown to induce malformations in zebrafish larvae [8]. In that report, larvae were exposed repeatedly for 6 days, and heart edema, curvature, and a reduced size of the swim bladder were observed at 25 µM. Liu et al. focused on cardiovascular toxicity of acetochlor and showed that acetochlor larval exposure causes pericardial edema, altered circulation, hemorrhage and thrombosis [9]. They determined mortality $LC_{50}$s ranging from 42-148 µM, depending on the length of exposure. Our results largely agree with these previous studies, both regarding the type of malformations observed and the micromolar range of concentrations required to cause malformations and mortality.

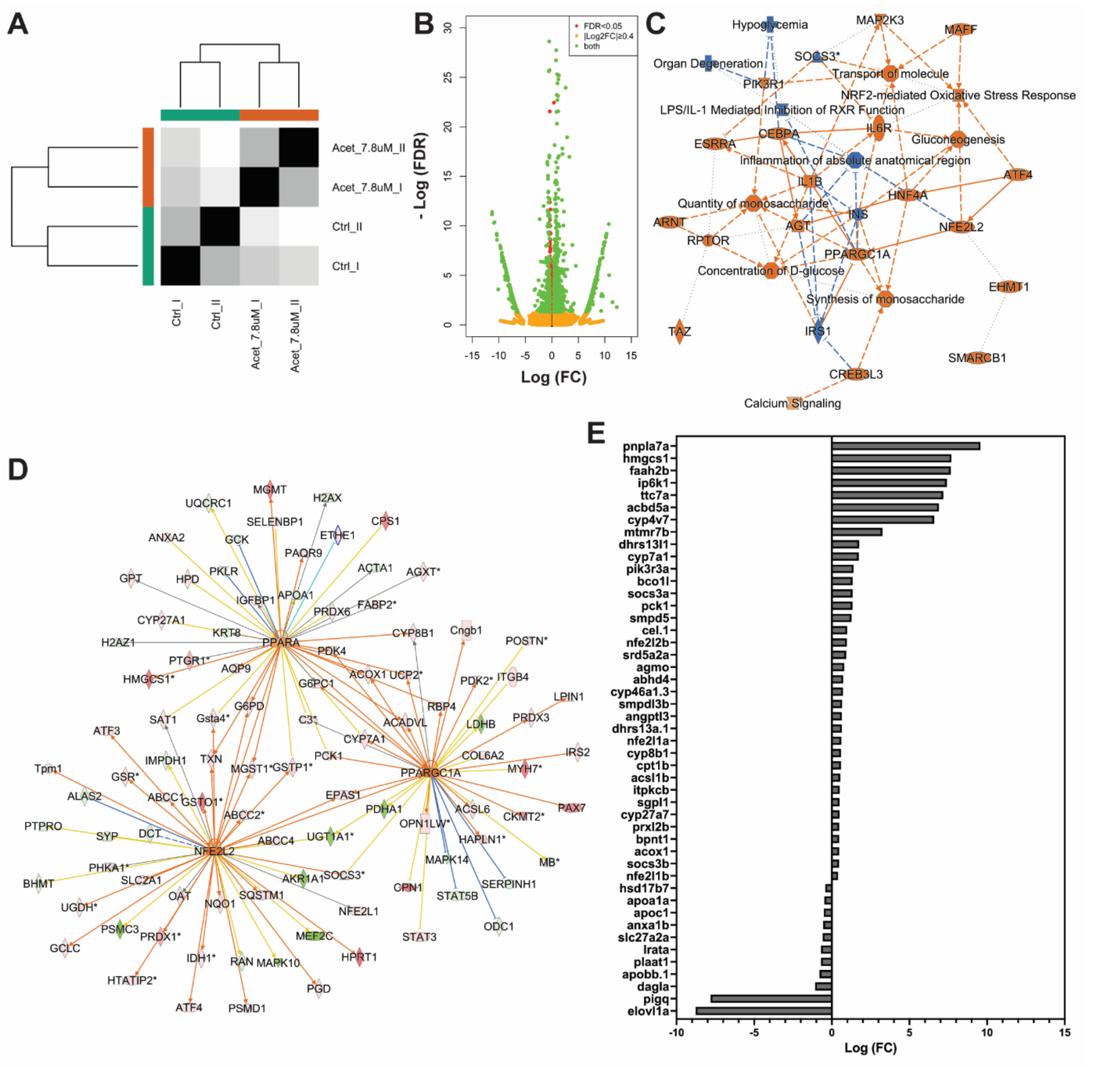


**Figure S6. RNA sequencing and pathway analysis of acetochlor treated zebrafish larvae.** At 48 hpf zebrafish larvae were treated with 7.8 µM acetochlor and RNA were collected at 72 hpf (24 hr exposure). **A)** Correlation between reads of mRNA-seq data of control and treated replicates. **B)** Volcano plot of log2 (fold change) and –log10 (p adjusted = FDR) of differentially expressed genes (DEG). False discovery rate (FDR) < of 0.05 and |log2 fold change| ≥ 0.4 were used as the cutoff criteria to select DEGs. 1211 genes were differentially expressed, 754 were upregulated and 457 downregulated. **C)** Graphical summary of the effected biological, molecular, and cellular functions showed an enrichment in many molecules that are involved in lipid and glucose metabolism. **D)** NFE2L2, PPARGC1A, and PPARA regulated genes. **E)** Genes involved in lipid metabolic pathway. The color intensity of the symbols in C and D represents the fold change of upregulated (red), downregulated (green). Orange represents predicated activation and blue represents predicated inhibition.

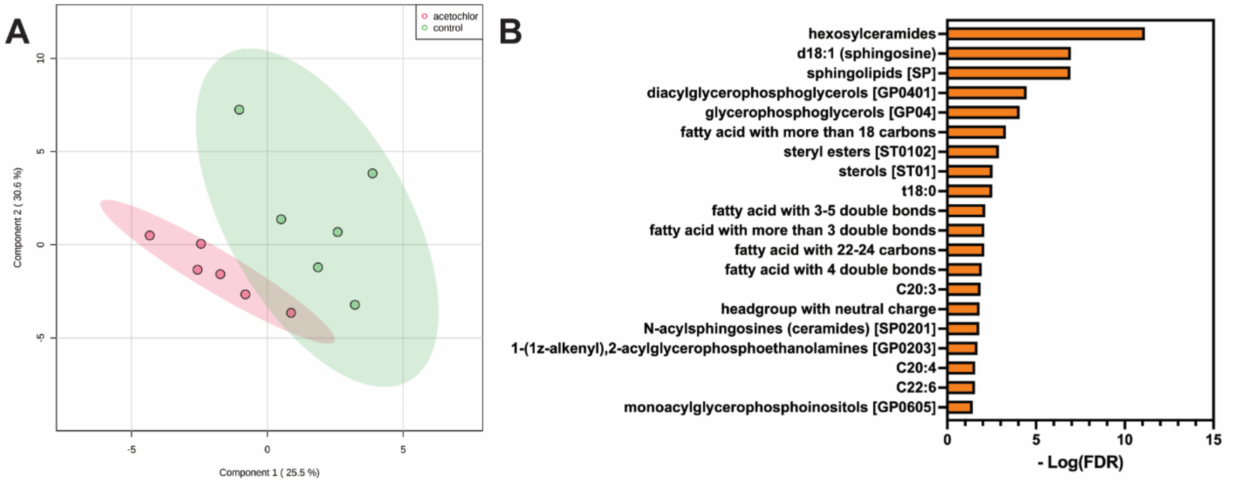


**Figure S7. Lipid signatures in zebrafish larvae treated with acetochlor revealed by lipidomics**. At 48 hpf zebrafish larvae were treated with 7.8 µM acetochlor and snap frozen at 72 hpf for lipidomic analysis. **A)** Partial least-squares discriminant analysis (PLS-DA) of lipid features in sample treated with vehicle control (green) or acetochlor (pink) of data obtained and merged from positive and negative electrospray ionization modes. **B)** LION (lipid ontology) enrichment analysis of exposed and control groups.

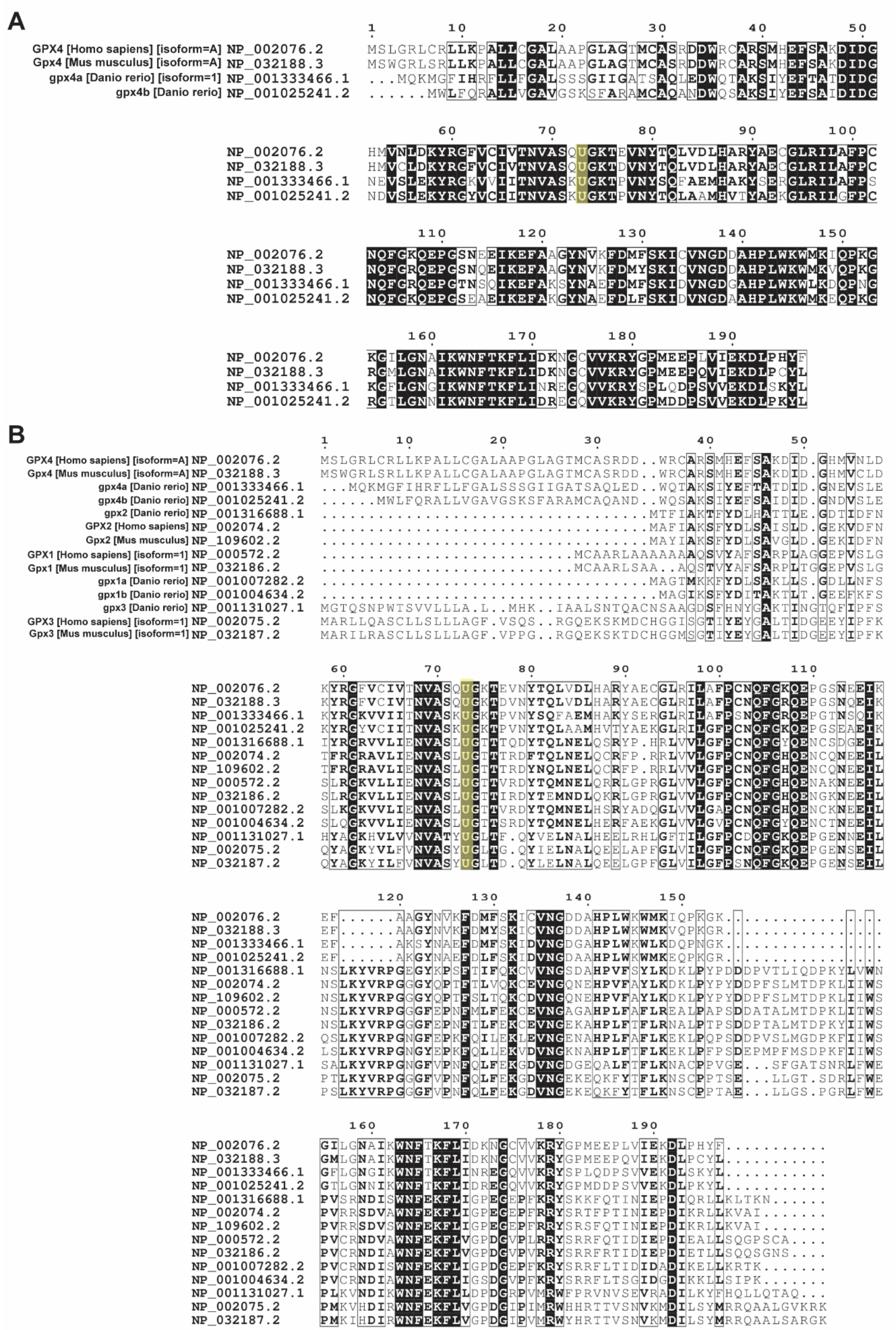
A
GPX4 [Homo sapiens] [isoform=A] NP_002076.2
Gpx4 [Mus musculus] [isoform=A] NP_032188.3
gpx4a [Danio rerio] [isoform=1] NP_001333466.1
gpx4b [Danio rerio] NP_001025241.2
B
GPX4 [Homo sapiens] [isoform=A] NP_002076.2
Gpx4 [Mus musculus] [isoform=A] NP_032188.3
gpx4a [Danio rerio] NP_001333466.1
gpx4b [Danio rerio] NP_001025241.2
gpx2 [Danio rerio] NP_001316688.1
GPX2 [Homo sapiens] NP_002074.2
Gpx2 [Mus musculus] NP_109602.2
GPX1 [Homo sapiens] [isoform=1] NP_000572.2
Gpx1 [Mus musculus] [isoform=1] NP_032186.2
gpx1a [Danio rerio] NP_001007282.2
gpx1b [Danio rerio] NP_001004634.2
gpx3 [Danio rerio] NP_001131027.1
GPX3 [Homo sapiens] [isoform=1] NP_002075.2
Gpx3 [Mus musculus] [isoform=1] NP_032187.2

**Figure S8. The selenocysteine of GPX4 is conserved through species and GPXs.** Amino acid sequence comparisons between GPX4 from human, mouse, and zebrafish **(A)** and GPXs 1, 2, 3 and 4 **(B)**. The conserved selenocysteine (U) is highlighted. Clustal Omega was used for multiple alignements and ESPript 3.x was used for viulaization. Other conserved amino acids are boxed in black color. Bold letters indicate peptide resulted from PROSITE search. Sequences were obtained from NCBI (https://www.ncbi.nlm.nih.gov).
